# *Elephants in a teacup*: Ranging and Habitat-use of Asian elephants *Elephas maximus* in a Plantation Dominated Landscape in Southern Western Ghats, India

**DOI:** 10.1101/2024.01.08.574633

**Authors:** Sreedhar Vijayakrishnan, Ganesh Raghunathan, Mavatur Ananda Kumar, Anindya Sinha

**Affiliations:** School of Natural Sciences and Engineering, National Institute of Advanced Studies, Bangalore, India; Nature Conservation Foundation, Mysore, India; Indian Institute of Science Education and Research, Kolkata, India

**Author notes:** Centre for Conservation and Research, Tissamaharama, Sri Lanka (Sreedhar Vijayakrishnan). Corresponding author (SV). **Author Contributions** **Conceptualisation:** Sreedhar Vijayakrishnan, Mavatur Ananda Kumar, Anindya Sinha **Data acquisition and curation:** Sreedhar Vijayakrishnan, Ganesh Raghunathan **Formal analysis:** Sreedhar Vijayakrishnan, Mavatur Ananda Kumar, Anindya Sinha **Funding acquisition:** Mavatur Ananda Kumar, Sreedhar Vijayakrishnan, Ganesh Raghunathan **Writing – original draft:** Sreedhar Vijayakrishnan, Anindya Sinha **Writing – review and editing:** Sreedhar Vijayakrishnan, Ganesh Raghunathan, Mavatur Ananda Kumar, Anindya Sinha.

## Abstract

Ranging patterns of four focal herds using a plantation-forest matrix of the Valparai plateau was studied by following the individuals on a regular basis, recording all possible sightings, for five years. Although surrounded by a network of protected areas, the focal herds were found to use the plateau extensively. There was minimal spatial overlap observed between the four herds, except between two, PTH and MON, which showed a fairly large overlap of 130 sq.km. The observed ranges of elephants were smaller (119.71 ± 24.11 sq.km) compared to previous studies on the species in India, but were comparable with those from Sri Lanka. Possible risk avoidance strategy is observed in their use of the landscape, as evidenced by the use of high intensity human activity areas mostly during night than day. Compared to previous studies from the same landscape, there is also an observed increase in the use of natural vegetation (in the form of rainforest fragments) than the plantation and monoculture habitats, indicating the importance of forest remnants for elephants on the plateau. The observations indicate importance of anthropogenic areas outside protected areas as potential habitats and, not just as movement pathways or as temporary refugia. Conservation planning should therefore consider such areas while initiating landscape-level management strategies for the species. The study also highlights the importance of longitudinal observation-based studies in ascertaining ranging, in the absence of logistically challenging techniques such as radio telemetry.

## Introduction

Increasing demand for natural resources combined with land-diversion for large-scale infrastructural projects, and agricultural expansion to meet its growing demand has contributed to the fragmentation of elephant habitats across Asia [1]. Furthermore, this has led to elephants ranging within and close to human-use landscapes (HUL), often resulting in intense human–elephant interface. Resulting escalation of human–elephant conflict lead to persecution of elephants in several HUL, and as a result threaten their populations. Persistence of elephants in human-use landscapes, hence, needs to be closely understood for their long-term conservation. This would include understanding their usage of such habitats, and their ranging and movement patterns through anthropogenic spaces.

It is estimated that, today, approximately 60 to 70% elephants in India range outside protected areas [2]. This is comparable with estimates from other range countries such as Sri Lanka, where roughly 70% of the geography is shared by elephants and people [3]. In countries like India, protected area sizes are also extremely small, and are often fragmented, resulting in elephant ranging in neighbouring human-use areas [4]. Elephants, as generalist species, use a wide range of habitats in natural, contiguous areas, with highest densities seen in deciduous habitats [5]. The general assumption continues to be that elephants use HUL as movement pathways or as temporary refuges, but recent studies indicate otherwise [3, 6]. Elephant ranging and distribution is often influenced by ecological factors, such as habitat types and vegetation [7, 8], and seasonal fluctuations in resource quality and quantity [9, 10]. In addition to this, behavioural factors also determine elephant ranges, in accordance to social dynamics [11, 12]. The anthropocene and its associated facets are additional aspects influencing elephant distribution, habitat-use and ranging.

Despite intense conflict, the cultural role that elephants have among communities across Asia has fostered tolerance for co-existence in shared landscapes, as is the case in India [13]. The country supports the largest wild Asian elephant population, which occurs in highly heterogeneous landscapes. Here, long-term conservation of elephants hinges on effective conflict resolution and concomitantly, increasing the overall resilience of the landscape to support elephant populations. Elephant home ranges often span many hundred square kilometres of mosaic landscape, which may include areas outside the reserves as well. The Protected Areas (PA) in India are often small (average reserve size: <300-km^2^) relative to elephant ranges and often are not delineated based on the species-specific requirements. Hence, PA-centric approach is acknowledged to be inadequate for elephant conservation, and the situation calls for an integrated landscape approach. This warrants fine-scale information about elephant populations, their ranging patterns, and their distribution across human-use spaces at range-country level. While within reserves where human imprint on the habitat is usually minimal, management priorities range from enhanced protection from poaching and habitat enrichment [14], in modified interfaces, or in anthropogenic landscapes with high zone of interaction between elephants and people, management priority is conflict resolution [3]. Understanding the dynamics of elephant movement patterns and habitat-use is thus crucial for this as well. Additionally, most studies aimed at understanding elephant ranging have been protected area centric in the past, and considering the species’ use of human-use areas, ascertaining their ranging and habitat-use in such landscapes becomes important.

The best approach to estimate elephant home ranges would be through telemetry approaches as has been done previously in Africa and Asia [8, 15-21]. However, logistical difficulties often challenge the application of this technique in many difficult habitats in Asia. And moreover, in some landscapes, following elephants regularly is relatively easy due to better visibility, aiding in direct data collection non-invasively.

This paper discusses the attempts at understanding fine-scale ranging patterns of four focal herds using the plantation-forest matrix of Valparai plateau in the Anamalai hills in south India, from movement data collected exclusively through direct observations. The objective of this chapter is to examine the home ranges of four focal herds in the study area, simultaneously also assessing their habitat-selectivity in the heterogeneous, modified, plantation-dominated landscape.

## Methods

The study was conducted on the Valparai plateau in the Anamalai hills, southern Western Ghats, which is a 220-km^2^ matrix of tea, coffee, and cardamom plantations interspersed with rainforest fragments and eucalyptus plantations (Fig 1). The landscape hosts about 70000 people [22] and supports an approximate 70 to 100 elephants that use the plateau regularly [7] Though seen throughout the year on the plateau, elephant movements peak between September and April, with sighting records dropping in the months of May to August. The four herds discussed in this paper, and few other peripheral groups with limited observations, are being tracked regularly as part of a long-term elephant-monitoring programme in the landscape. For ranging and habitat selectivity, data from four of those five herds were analysed.

**Figure 1:**
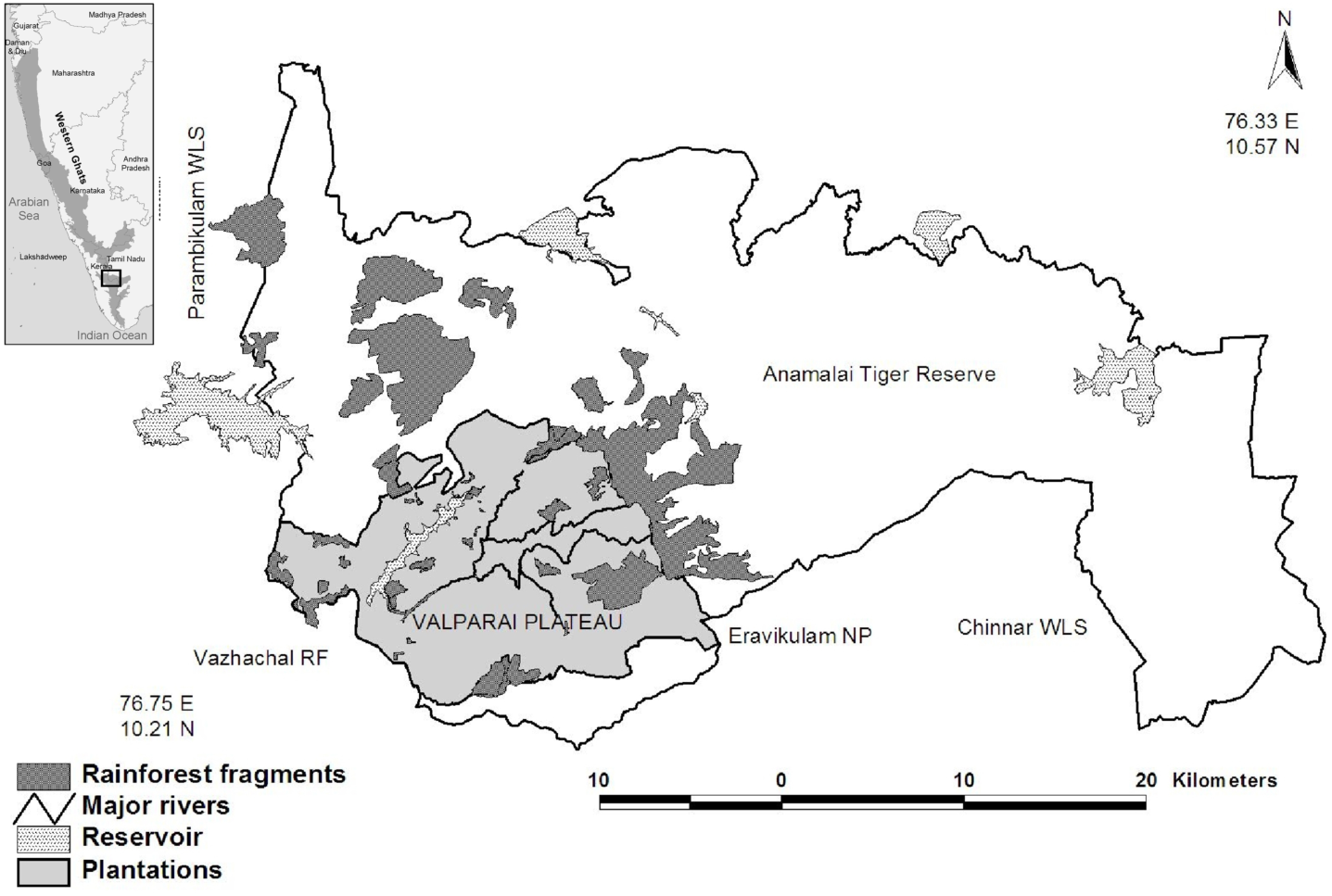
Study area.

### Elephant Tracking

The tracking of elephants was carried out as part of the long-term elephant monitoring programme on the plateau, with primary focus on understanding their movement patterns to aid conflict mitigation. Tracking team follows regular and a few peripheral herds within plantations, recording information on herd ID, number of individuals sighted, GPS location, type of habitat, time, date, and conflict incidents if any. Elephants are usually tracked on foot, and record GPS coordinates using a handheld Garmin eTrex 30 GPS during direct observations and noting indirect signs such as fresh dung, feeding, footsteps of sightings at regular intervals along the path of movements.

### Data extraction and analysis

To deal with the variation in terms of number of data points per herd per day, primarily owing to the uncertainties in their daily sightings, one day and night data points each were chosen at random per herd and per date, using the statistical program R [23]. These data points were further used for estimating home ranges and habitat-use analyses. Tracking data collected between 2013 and 2017 (five-year period) were used for the home range analyses of focal herds. For analysing the habitat-use patterns of all elephants using the plateau, data collected from 2013 January till 2019 March was used. The last two years of data were not used for home-range analyses owing to difficulties in assigning systematic herd-level ID to data points on many occasions by the tracking team. Habitat-use patterns were also estimated using package adehabitat [24] on program R [23], using Manly’s selectivity index.

Program adehabitat was used for assessing home ranges and intense use areas in minimum convex polygon (MCP) and kernel density estimator (KDE) frameworks. 50%, 75%, and 95% KDE were estimated for all four herds using the program. Home range visualisation and creation of maps, and calculating home range sizes and overlap areas were performed on open source spatial analytical program QGIS ver. 3.10 [25].

## Results

### Home range estimates of focal herds –– MCP

A total of 1816 data points collected from four focal herds over five years were used for MCP analysis. Initial observations indicate that the study herds, as evident from Fig. 2, extensively use the plateau, covering most of it geographically. The home ranges (100% MCP) ranged from 71.37 sq. km (STP group) to 165.53 sq.km (MON group). While there is visible spatial segregation in terms of the area of usage, there is significant overlap between the MON and PTH groups. With 165.53 sq.km and 156.60 sq.km respectively, the MON and PTH groups use the North and North-western parts of the plateau predominantly, with an overlap of 129.4 sq.km. MON group uses the coffee, Eucalyptus and the intense human–use areas of the landscape, compared to the PTH group that uses the riparian more. All four groups invariably use the Nadu Ar-Sholayar riverine vegetation, which is discussed in a later section in this chapter. The PCH (85.37 sq.km) and STP (71.37 sq.km) groups using southern side of the plateau showed a minimal overlap of 15.22 sq.km.

**Figure 2:**
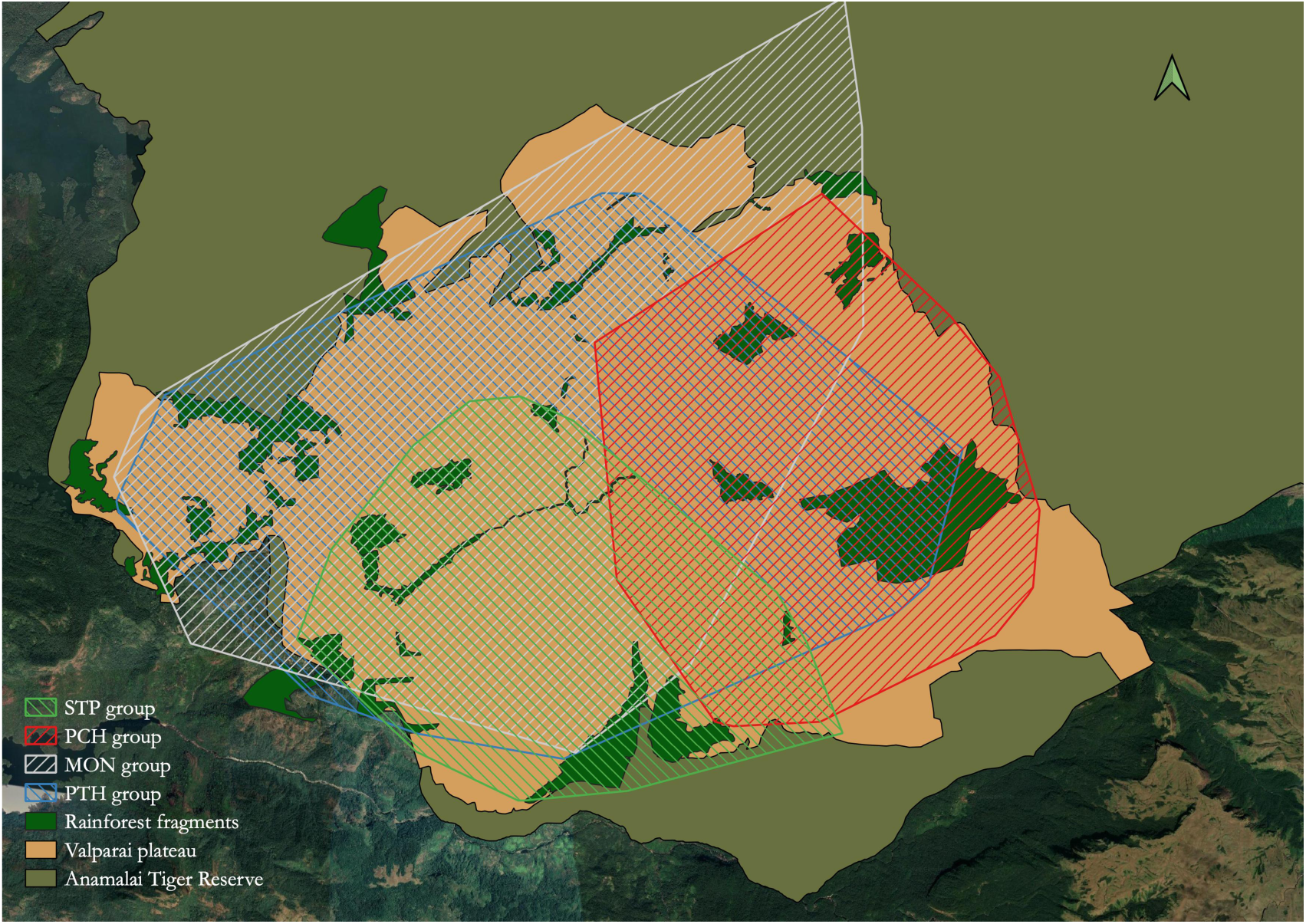
Minimum Convex Polygons (100%) showing home-range of all four study herds on the Valparai plateau.

### Intense use areas –– KDE

While the herds seemingly had large ranges on the plateau, their intense use areas were restricted to smaller areas, as understood through Kernel Density Estimation. Intense use areas of the herds are as in table 1. The STP group with the smallest KDE uses the tea plantations in the southern part of the plateau, adjoining the neighbouring Malayattoor division, and the PCH with the next smallest range uses tea plantations fragments in the East and South-eastern parts of the landscape.

**Table 1:**
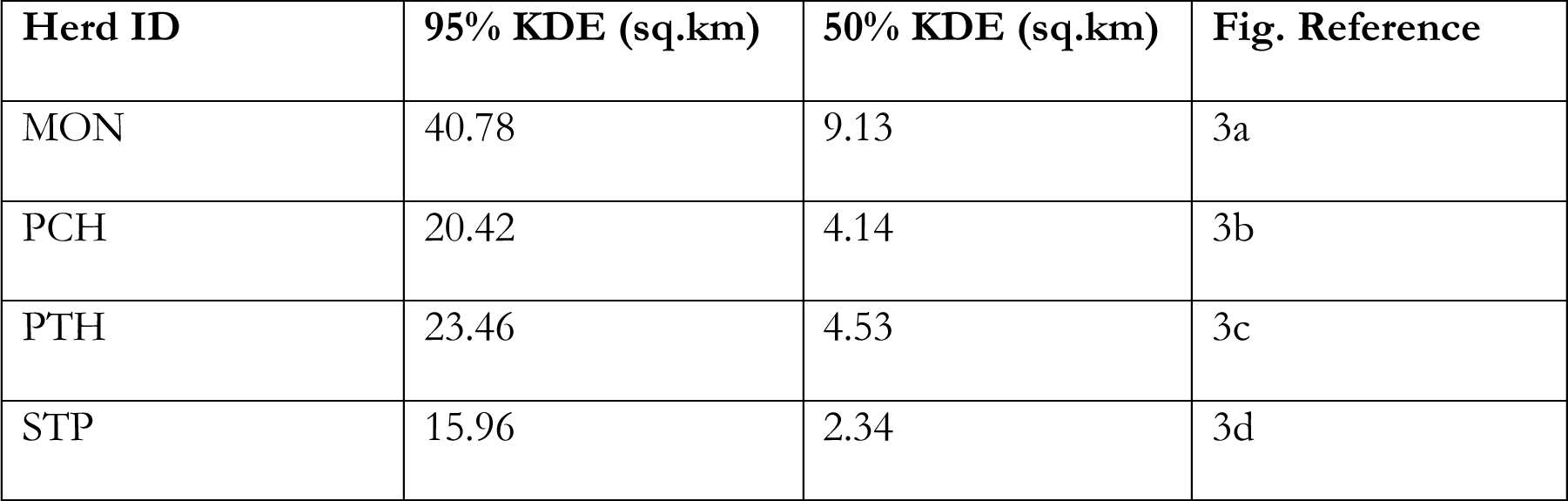
95 and 50% KDE of the four study herds, showing their intense use areas.

**Figure 3a:**
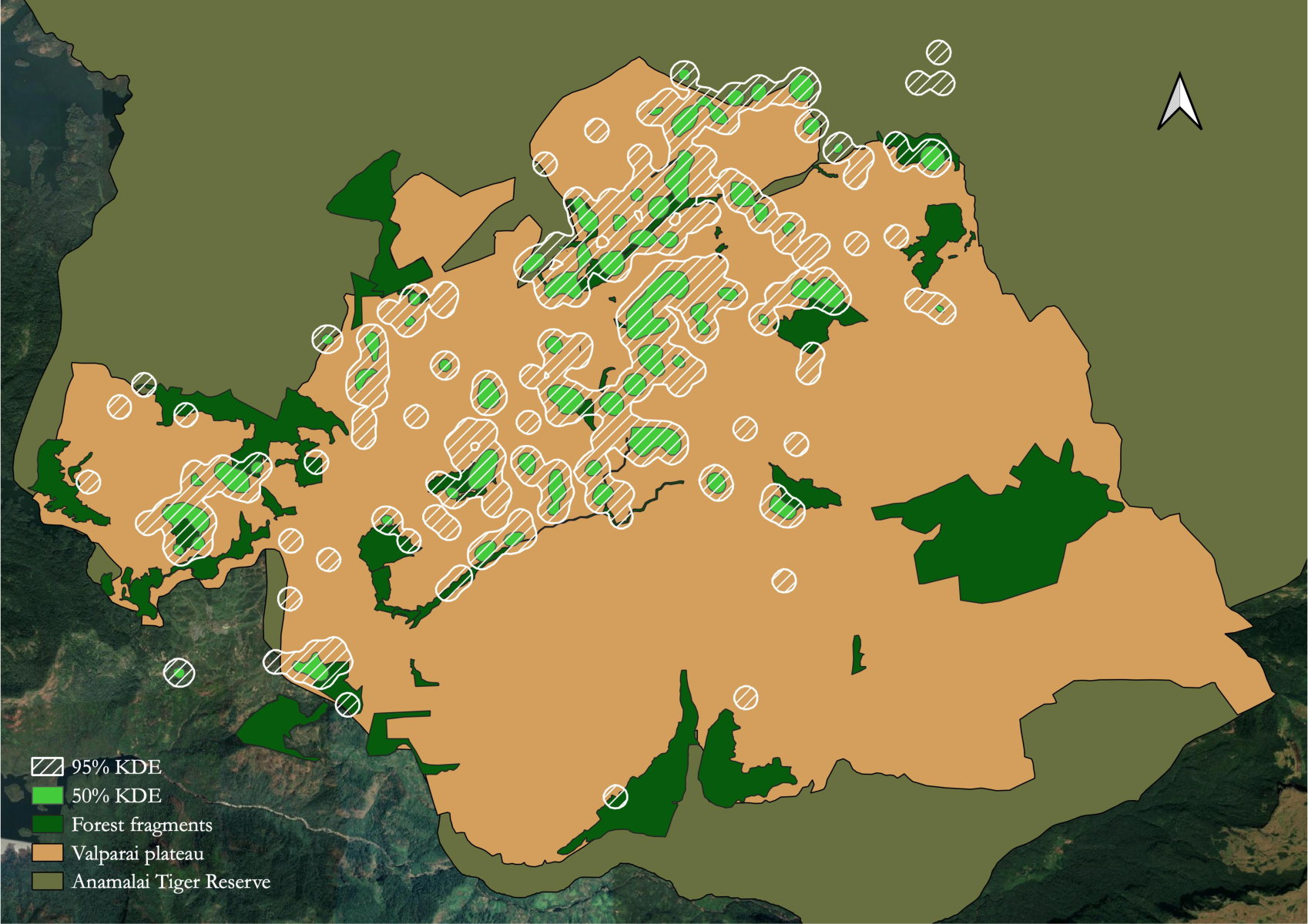
Core use areas (95% and 50%) of MON herd.

**Figure 3b:**
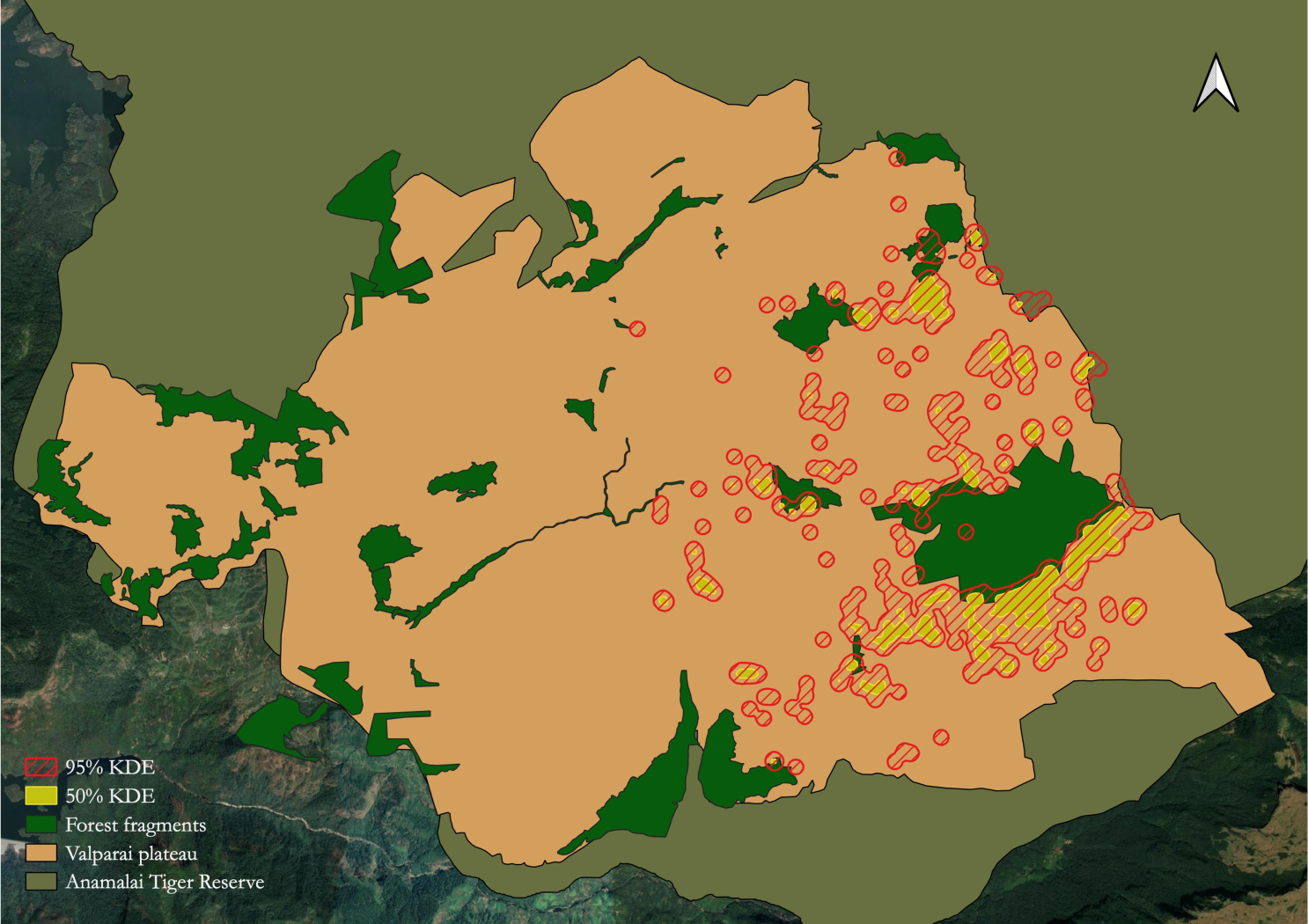
Core use areas (95% and 50%) of PCH herd.

**Figure 3c:**
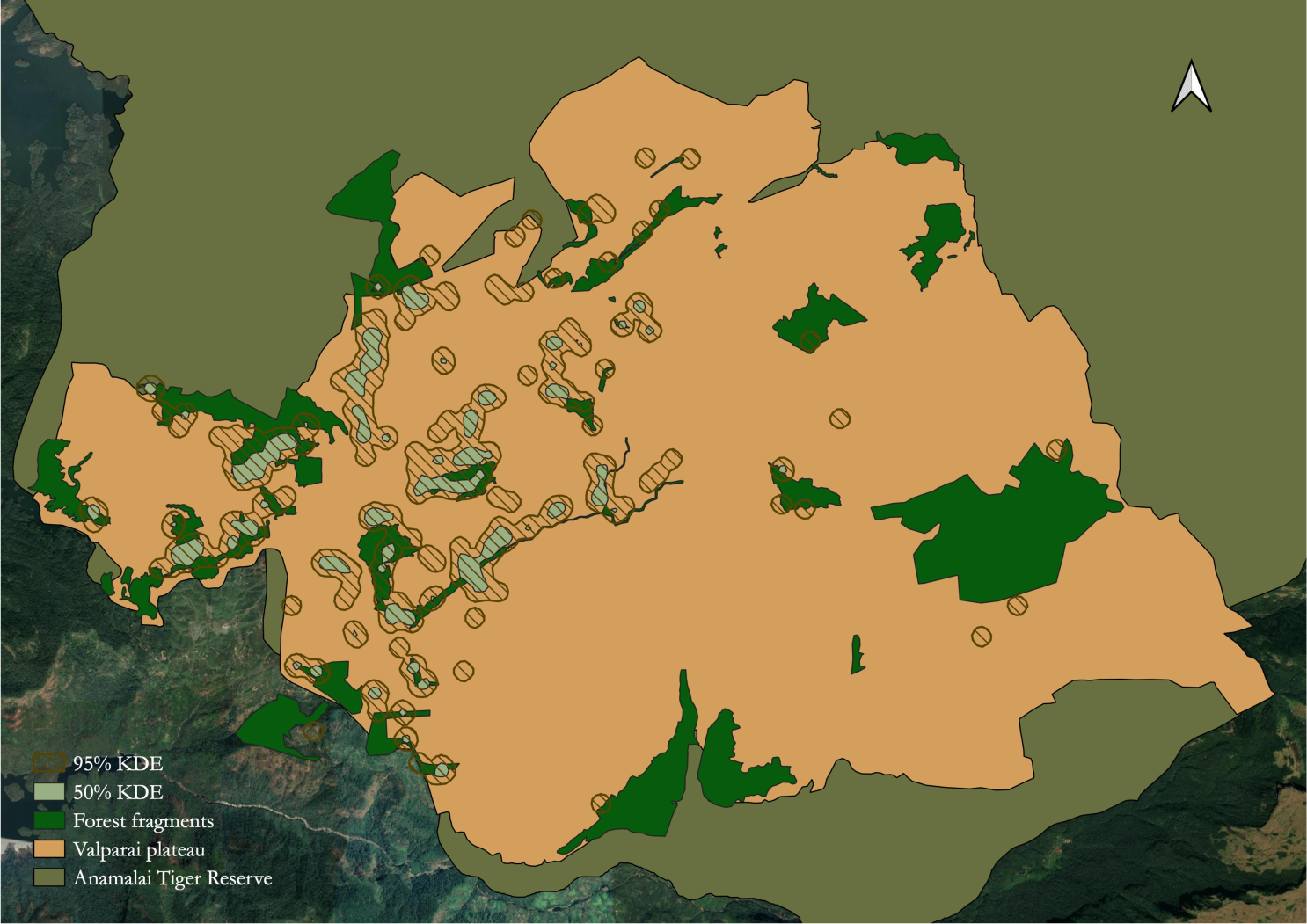
Core use areas (95% and 50%) of PTH herd.

**Figure 3d:**
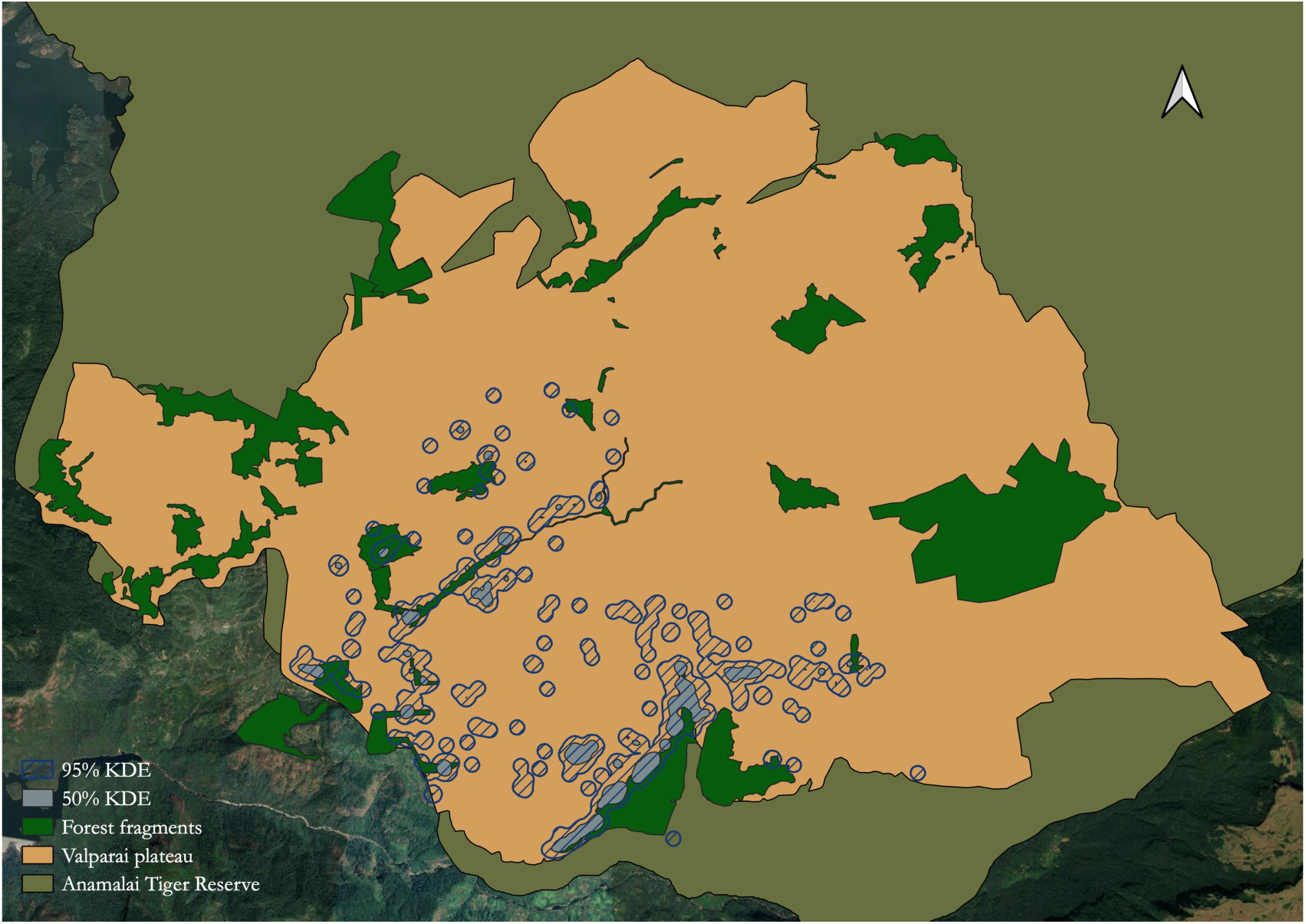
Core use areas (95% and 50%) of STP herd.

### Habitat-selection patterns in the plantation-dominated mosaic

A total of 2728 data points between 2013 and 2019 were used for assessing habitat selectivity by elephants using Manly’s selectivity index. There was a significantly high selection of riparian habitats (Manly’s selectivity index, *w_i_*= 59.325, SE = 4.52, p < 0.001; Fig. 4, followed by natural habitats (Manly’s selectivity index, *w_i_*= 11.858, SE = 0.228, p < 0.001). Tea (Manly’s selectivity index, *w_i_*= 0.392, SE = 0.013, p < 0.001) and other (which includes rocky outcrops, grasslands etc.) habitat types (Manly’s selectivity index, *w_i_*= 0.147, SE = 0.062, p < 0.001) were, however, less selected.

**Figure 4:**
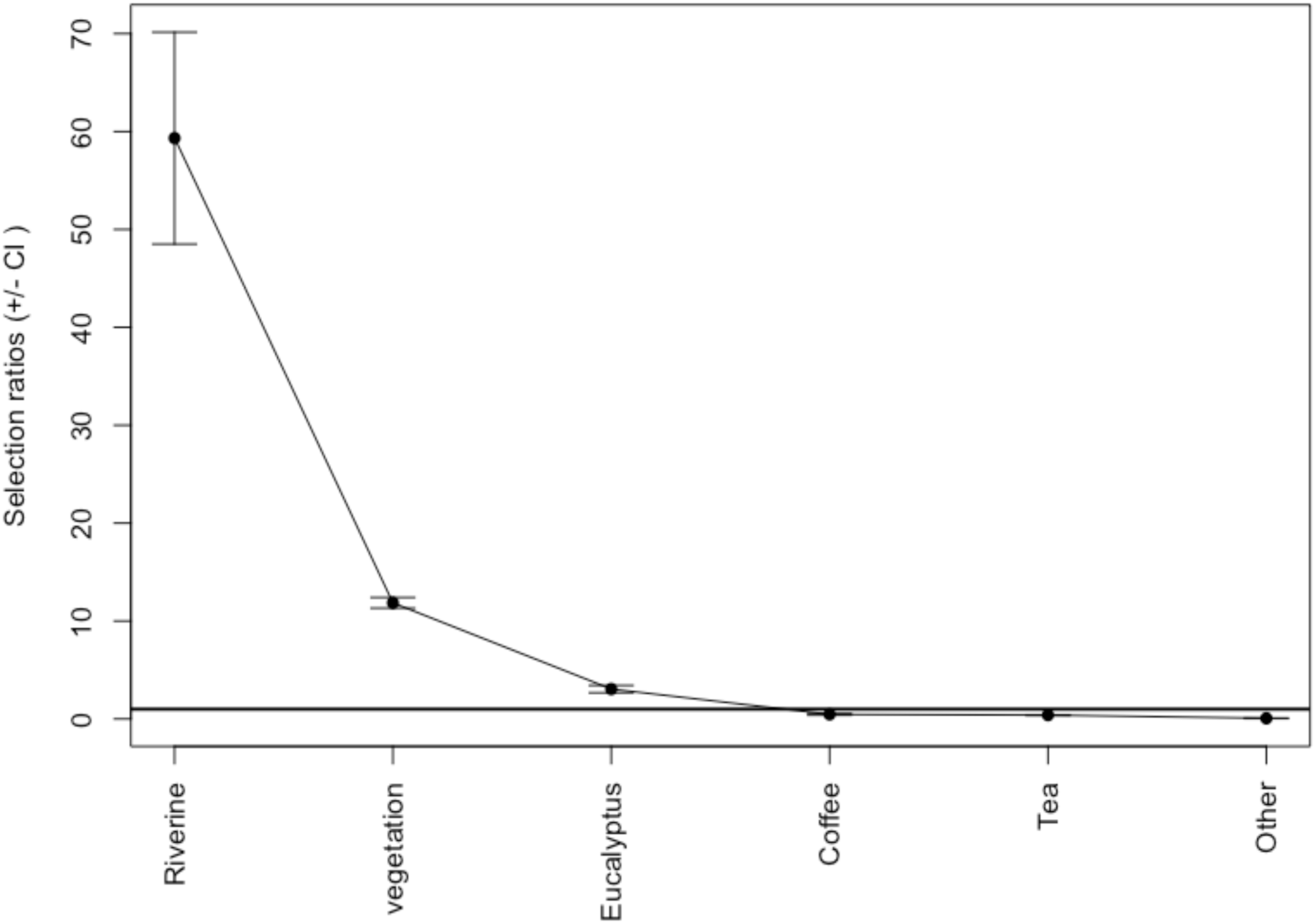
Overall Manly selectivity index of study herds on the Valparai plateau.

Tea habitats were used less during day hours (Manly’s selectivity index, *w_i_*= 0.171, SE = 0.011, p < 0.001; Fig. 5a) than during night hours (Manly’s selectivity index, *w_i_*= 0.857, SE = 0.026, p < 0.001; Fig. 5b). Daytime selectivity was more towards riparian habitats (Manly’s selectivity index, *w_i_*= 70.664, SE = 5.952, p < 0.001), followed by natural vegetation (Manly’s selectivity index, *w_i_*= 14.487, SE = 0.27, p < 0.001), and Eucalyptus (Manly’s selectivity index, *w_i_*= 3.803, SE = 0.209, p < 0.001). There, however, was a decline in the selection of these habitats during night hours, with a subsequent increase in that of tea and coffee habitats.

**Figure 5a:**
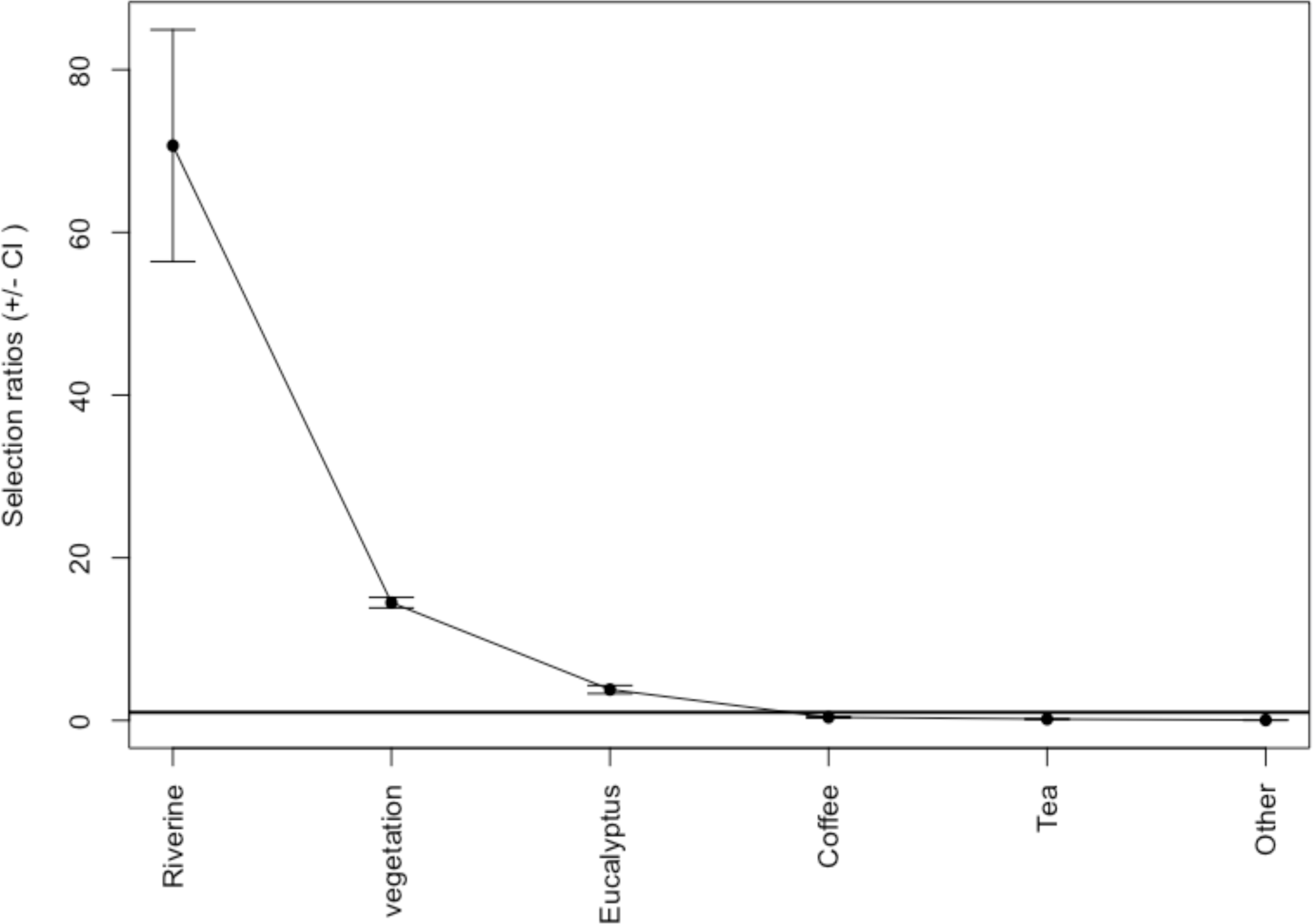
Diurnal Manly selectivity index of the study animals.

**Figure 5b:**
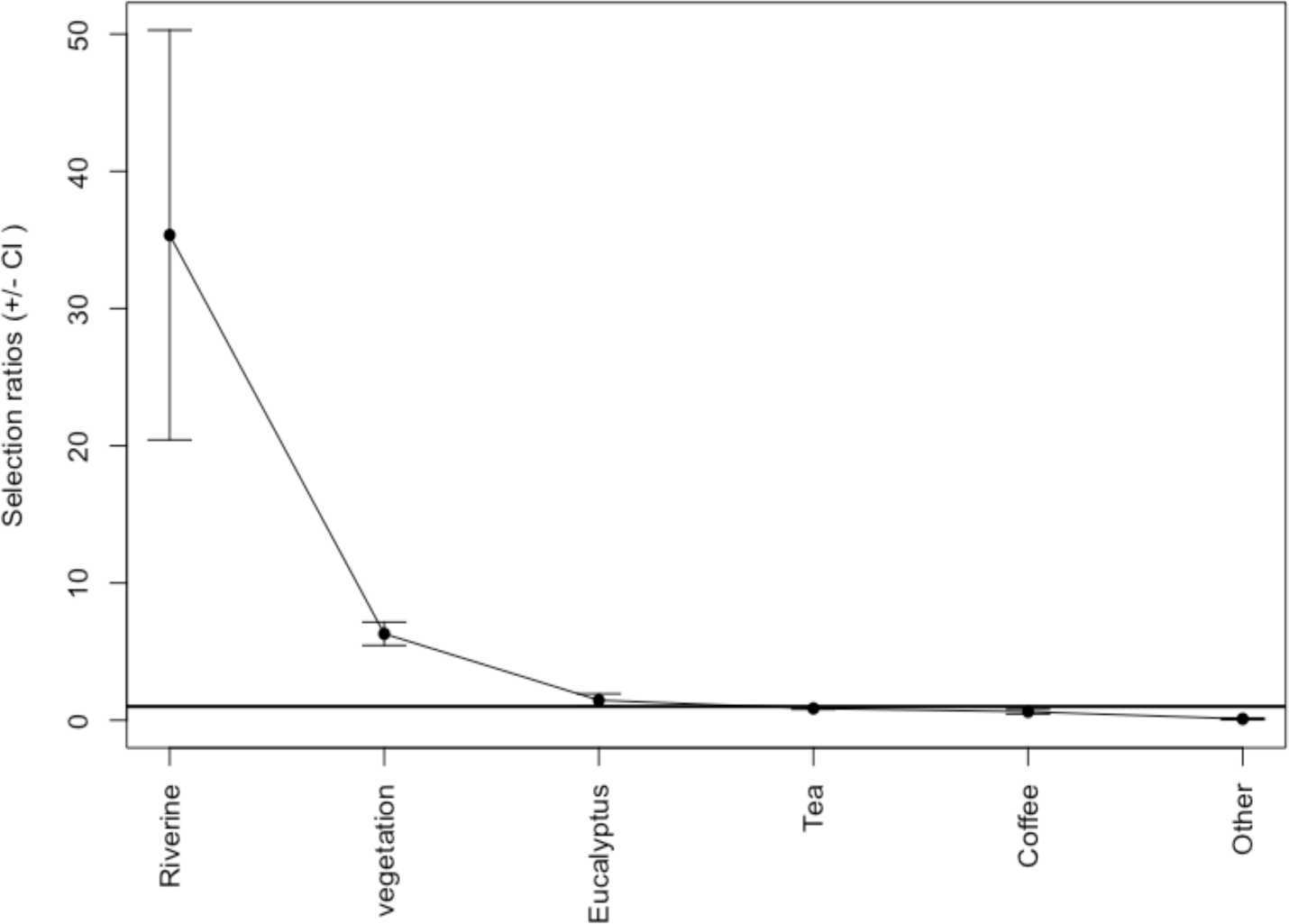
Nocturnal Manly selectivity index of the study animals.

**Table 2:** Season-wise frequency of elephant locations in different habitat types on Valparai plateau.

| Habitat | Dry |  | Wet |  |
| --- | --- | --- | --- | --- |
|  | Expected | Observed | Expected | Observed |
| Coffee | 63.92 | 80 | 104.08 | 88 |
| Eucalyptus | 126.71 | 129 | 206.29 | 204 |
| Natural Vegetation | 517.48 | 513 | 842.52 | 847 |
| Other | 9.51 | 13 | 15.49 | 12 |
| Riverine | 61.64 | 90 | 100.36 | 72 |
| Tea | 258.74 | 213 | 421.26 | 467 |

## Discussion

### Ranging in a human-dominated landscape

The patterns observed in this study based on data compiled across five years indicate that the study individuals have home ranges comparable with earlier estimates [20, 26]. However, the core use areas of the groups discussed in this study are extremely small. This could be related to various factors ranging from resource availability, disturbance factors or intensity of human activity, and inter-herd social dynamics [27]. Consistent availability of forage with minimal disturbances could contribute to small and stable home ranges. In disturbed habitats with intense anthropogenic pressures, ranges often tend to be as big as a few thousand square kilometres, as seen in landscapes such as Nilgiris, Eastern Ghats, or North-Central India [28, 29]. Earlier studies have shown range shifts in elephants in response to human disturbances and conflicts [29]. However, little is known about elephant groups that exclusively use HUL and range within disturbed or modified landscapes. Overall movement patterns of the focal herds also indicate home range fidelity seemingly exhibited by the focal herds, which has previously been recorded by an earlier study in this landscape [30], and in Sri Lanka [20].

The home range sizes of the focal groups seem to negatively correlate with group sizes, with the smallest group (MON group; 5 individuals at the time of the study, now 6) having the highest home range, and the two large groups (STP and PCH groups with 13 and 16 individuals respectively) having smaller home ranges. This is reflected in their overall ranges and core use areas. This is, however, in contrary to the expansionism concept explained by Macdonald [31] in carnivores where increase in pride biomass results in increase or expansion of home ranges [32]. In a human-dominated landscape such as the study area, the chances of being detected are fairly high, owing to spatial overlap between elephants and humans. This further means increased exposure to stressors for the individuals. Small groups perhaps have lower probabilities of being detected, and are better able to navigate through the anthropogenic spaces and associated threats. The groups with smaller home ranges also have the highest number of size class 1 and 2 juveniles. This could possibly be because of the fact that small, dependent juveniles could possibly add to inherent risks faced by the herd, and in turn affect overall behavioural patterns [33]. This further influences their movement patterns, wherein groups would adopt risk-averse strategies and range in low-disturbance areas [34], contiguous with the PA.

The general assumption remains that large mammals such as elephants tend to use HUL as temporary refuges or as movement pathways. Behavioural modifications enable elephants to use refugia in human-use areas as a strategy to overcome the impacts of fragmentation [35]. However, our five-year data shows otherwise, that with minimal drop in sighting frequencies in wet season, elephants, especially the focal groups have been observed to be using the plateau throughout the year. Vijayakrishnan et al [36] also indicate that the study individuals could also potentially be exhibiting some level of adaptation to living in a human-dominated landscape. A large part of their range seems to be within the plateau, as is described in the earlier study in the landscape [7], and it seems unlikely that they would be ranging much larger area outside the plateau during the wet season. In addition, elephants are known to use heterogeneous habitats extensively, considering the high level of nutrition available in such areas [20, 37].

### Temporal differences in habitat use by elephants

In corroboration with findings of an earlier study in the same landscape [7], this study showed that tea habitats were used predominantly during night hours for movements between natural vegetation habitats and possibly as a risk avoidance strategy. Daytime use of riparian and other natural vegetation reiterates the importance of diurnal cover in terms of resource and shelter requirements in risk-prone areas. This is similar to the findings on African elephants, where animals spent more time at night in human-use landscapes, especially in risk-prone areas [35]. The use of riverine area more during dry season also indicates the importance of water, especially natural sources in defining elephant habitat-selectivity patterns, as explained in earlier studies [38, 39].

Factors ranging from resource availability [40], human activity [18], and bioclimatic factors affect seasonality in elephant ranging and habitat-use, leading to switch between browse and graze [5]. There, however, may not be such pronounced variation in evergreen habitats. In this study, seasonal changes were primarily observed in riverine and tea habitats. Seasonal variation in habitat-use shows preference for tea in wet season, which could possibly be due to the availability of grasses in swamps amidst tea estates. Kumar et al. [7] showed how the riparian segments in the study area are extensively used by the species, indicating their crucial role in a habitat-matrix such as this. This finding is now reiterated in the current study, wherein the riparian vegetation continues to be disproportionately used by elephants. This is in further corroboration with earlier observations that riverine habitats are crucial habitats for large mammals such as elephants 40. In addition, there is also an observed increase in the focal groups’ usage of the remnant forest fragments. This could possibly be a result of the ongoing restoration efforts that are attempted at rejuvenating these areas [41]. This also underlines the importance of habitat restoration programmes for long-term species conservation. Due to the presence of humans, elephants largely avoided other habitats such as roads and residential colonies.

### Observed herd-level changes in ranging patterns

A previous study on the STP group ranging patterns identified an area of 114 sq.km being used by the individuals [30]. The group, which used to be 19 individuals split into two subgroups as STP group and PCH group in 2011-12, and have been observed as two distinct social units since then, with extremely minimal (n<5) observed direct interactions. The group ranging patterns have also accordingly changed, splitting into two distinct ranging areas with an overlap of about 15 sq.km. The long-term monitoring in this case helped in recording this split, and subsequent changes in their ranges, perhaps previously unrecorded from Asian elephants. Similarly, there is also an observed shift in change in ranging patterns of the PTH group, evidenced by extension of their previously known area, after PT, the older female in the group dispersed. This observation warrants closer monitoring of the group to understand any potential role of matriarchal dispersal in expanding or shifting home ranges.

Closer observations of this kind would help us in understanding herd dynamics and its implications on their ranging patterns, especially in modified landscapes such as these. Although Fernando et al. [20] discusses the limitations of observational studies in determining home range sizes, our study shows that longitudinal monitoring could possibly help in understanding elephant ranging patterns in select landscapes.

## Acknowledgements

The authors wish to acknowledge the Tamil Nadu Forest Department and the Anamalai Planters Association for granting necessary permissions in carrying out this work. The authors also thank friends and colleagues at the Nature Conservation Foundation for their support and encouragement. Thanks are also due to Akshay Surendra and Rohit Jha for discussions with regard to the HR analysis.

